# Persistence of *Fusarium oxysporum* f. sp. *fragariae* in soil through asymptomatic colonization of rotation crops

**DOI:** 10.1101/464099

**Authors:** Peter M. Henry, Ana M. Pastrana, Johan H.J. Leveau, Thomas R. Gordon

**Affiliations:** Department of Plant Pathology, University of California, Davis 95616

## Abstract

Asymptomatic plant colonization is hypothesized to enhance persistence of pathogenic forms of *Fusarium oxysporum* in the absence of a susceptible host. However, a correlation between pathogen populations on living plant tissues and soilborne populations after tillage has not been demonstrated. Living and dead tissues of broccoli, lettuce, spinach, wheat, cilantro, raspberry, and strawberry plants inoculated with *Fusarium oxysporum* f. sp. *fragariae* (the cause of Fusarium wilt of strawberry) were assayed to quantify the incidence of infection and extent of colonization by this pathogen. All crops could be infected by *F. oxysporum* f. sp*. fragariae*, but the extent of colonization varied between plant species. Pathogen population densities on non-living crown tissues incorporated into the soil matrix were typically greater than those observed on living tissues. Crop-dependent differences in the inoculum density of *F. oxysporum* f. sp*. fragariae* in soil were only observed after decomposition of crop residue. Forty-four weeks after plants were incorporated into the soil, *F. oxysporum* f. sp. *fragariae* soil population densities were positively correlated with population densities on plant tissue fragments recovered at the same timepoint. Results indicate that asymptomatic colonization can have a significant, long-term impact on soilborne populations of Fusarium wilt pathogens. Cultural practices, such as crop rotation, should be leveraged to favor pathogen population decline by planting hosts that do not support extensive population growth on living or decomposing tissues.

## INTRODUCTION

Crop rotation can reduce severity and incidence of wilt diseases caused by *Fusarium oxysporum* (Fang et al. 2011; Gatch and du Toit 2017; Hopkins and Elmstrom 1984; Rao et al. 2014; Toledo-Souza et al. 2012; Wang et al. 2015). However, not all crop rotations lead to reduced disease pressure (Fang et al. 2011; Hopkins and Elmstrom 1984; Rao et al. 2014; Toledo-Souza et al. 2012; Wang et al. 2015). Soil populations of Fusarium wilt pathogens are likely to decrease substantially within 1 to 3 years under fallow conditions (Banihashemi and Dezeeuw 1973; Elmer and Lacy 1987; Gordon and Koike 2015; Rishbeth 1955; Scott et al. 2012; Stover 1956; Vakalounakis and Chialkas 2004), but a very broad range of plant hosts can be colonized asymptomatically by pathogenic forms of *Fusarium oxysporum* (Gordon et al. 1989; Haware and Nene 1982; Scott et al. 2014). Some plants support relatively high populations of Fusarium wilt pathogens and have a greater potential to increase inoculum (Dhingra and Coelho Netto 2001). It is hypothesized that differences in the efficacy of crop rotations are explained by corresponding differences in plant tissue colonization (Gordon 2017).

The relative risk of rotation with a particular crop is frequently estimated by quantifying pathogen biomass in living plant tissues (Dhingra and Coelho Netto 2001; Haware and Nene 1982; Leoni et al. 2013; Pastrana et al. 2017a; Scott et al. 2014). The extent of colonization of senescent or non-living tissue is less commonly examined. *Fusarium oxysporum* f. sp. *melonis* populations on the roots of seven crops were found to significantly increase within five days after plants were severed at the soil line (Gordon and Okamoto 1990). To the extent that pathogen growth occurs in senescing or decomposing tissue, colonization of living tissues may not indicate the true potential for a crop to increase soilborne inoculum. A combination of experiments that quantify inoculum densities on living and decomposed plant tissues, and a comparison of these values with soil populations after plant tissue incorporation, would help to establish protocols for identifying crops that support increases in populations of Fusarium wilt pathogens. Such experiments could test the implicit hypothesis that inoculum densities on living plant tissues are predictive of soilborne inoculum after tillage.

Fusarium wilt of strawberry, caused by *F. oxysporum* f. sp*. fragariae,* is a disease of serious concern for strawberry production regions worldwide, including those in California. Fusarium wilt resistant cultivars are available but yield losses may still occur in highly infested fields (Mark Bolda, personal communication). Furthermore, the best available fumigation practices can still result in 10 to 15% incidence of Fusarium wilt in the following season (Mark Bolda, personal communication; Koike and Gordon 2015). Both conventional and organic strawberry growers in California have shown interest in using crop rotation for disease control (Lloyd 2015), but the potential for crops commonly grown in California strawberry production regions to become colonized by *F. oxysporum* f. sp*. fragariae* remains unexplored. Therefore, experimental validation is required to ensure that crop rotations do not facilitate persistence of this pathogen in soil.

The objectives of this study were to: 1) assess the population density of *F. oxysporum* f. sp*. fragariae* on four types of living tissues of diverse hosts (including fine roots, transport roots, taproot/crown cortex, and taproot/crown stele), 2) determine if pathogen population density in host tissues increases after incorporating these tissues into the soil, and 3) evaluate the correlation between population density on decomposing host tissues and persistence in soil over time.

## MATERIALS AND METHODS

### Fungal transformation

To facilitate detection in plant tissues and soil, a known pathogenic isolate of *F. oxysporum* f. sp. *fragariae* (GL1080) (Henry et al. 2017) was transformed to obtain an antibiotic-selectable strain using a protocol described by Pareek et al. (2015). *Agrobacterium tumefaciens* strain AGL-1 containing the pBHt2 vector (Mullins et al. 2001) (kindly provided by Dr. Richard Bostock, Department of Plant Pathology, UC Davis) was grown on an LB agar plate supplemented with kanamycin (50 μg/mL) at 28°C for 2 days in darkness. LB broth supplemented with kanamycin (50 μg/mL) and rifampicin (50 μg/mL) was inoculated with a single colony and incubated for 20 hours at 28°C on a rotary shaker at 250 rpm. Cells were centrifuged, washed twice with sterile deionized water, resuspended to an OD600 of 0.15 in induction medium containing 200 μM acetosyringone (Bundock et al. 1995), and incubated for 4 hours at 28°C, 250 rpm.

The optimum concentration of hygromycin B and cefotaxime for transformation was determined as described by Yuan et al. (2014). GL1080 was cultured on potato dextrose agar (PDA) for 10 days at room temperature. Spores were scraped from the surface of culture plates by adding sterile deionized water and scraping gently with a sterile glass rod. The resulting suspension was filtered through two-layers of sterile cheesecloth and the filtrate was centrifuged at 2000 × *g* and 4°C for 10 min. The pellet was washed twice with sterile deionized water and diluted to a density of 10^6^ spores/mL.

The suspension of *F. oxysporum* f. sp*. fragariae* spores was mixed in a 1:1 ratio with acetosyringone-induced *Agrobacterium* culture. Two-hundred microliters of the combined suspensions were spread on nitrocellulose membranes (Whatman®, Cat. No. 7141-104) and placed on the agar surface of induction medium containing 200 μM acetosyringone. Other nitrocellulose membranes were inoculated with only the fungal spore suspension (no *Agrobacterium*) as a negative control. Plates were incubated at 28°C for 2 days, after which the membrane was transferred to *F. oxysporum* minimal medium agar plates (Correll et al. 1987) containing hygromycin B (100 mg/L) and cefotaxime (300 mg/L). Hygromycin B resistant fungal colonies appeared after 7 days of incubation at 25°C. No colonies were observed growing from the negative control membranes.

To check for phenotypic stability, transformed colonies were serially transferred five times on PDA plates containing hygromycin B (100 mg/L) and cefotaxime (300 mg/L). DNA was isolated from transformants as described by Pastrana et al. (2017b). To confirm the integration of the hygromycin B phosphotransferase gene (*hph*), an approximately 1-kilobasepair fragment was PCR amplified using primers HPH-F and HPH-R (Kemppainen et al. 2005). Transformants were single hyphal tipped and stored on dried filter paper (Gordon and Okamoto 1991). One of these transformants (GL1080-K) was used in subsequent experiments.

### Root tip colonization in naturally infested field soil

The substrate used in these experiments was a 3:2 mixture of naturally *F. oxysporum* f. sp. *fragariae*-infested field soil and autoclaved fine white sand (Capitol Sand and Gravel). Field soil was obtained in August of 2015 from a commercial field in Ventura County with a history of Fusarium wilt of strawberry, and all isolates from these soils were in the same somatic compatibility group as GL1080-K (Henry et al. 2017). The soil/sand mixture was prepared just prior to the start of each experiment (repeated 3 times).

Inoculum density of *F. oxysporum* f. sp. *fragariae* in the soil/sand mixture was estimated using soil dilution plating (Henry et al. 2017), followed by a PCR test of putative pathogen colonies using primers described by Suga et al. (2013). Previous experiments showed that 96.8% of *F. oxysporum* f. sp. *fragariae* isolates from this soil could be identified by the FofraF/FofraR primer pair (Henry et al. 2017). DNA was extracted by methods described in Henry et al. (2017). PCR verification was conducted as per Suga et al. (2013) with the following modifications: the PCR reaction mixture (25 μL) contained 1x GoTaq Green Master Mix, 0.4 μM FofraF and FofraR primers, and 40-100 ng/μL gDNA. Two technical replicates of the soil assay were conducted at the beginning of each experiment. Results from these assays were averaged to determine the mean abundance of *F. oxysporum* f. sp. *fragariae* in the soil of each experiment.

The soil/sand mix was dispensed into thirty-two 2 L pots, into which 14-day-old seedlings/transplants of the general rotation crop panel (Table 1) were planted. Plants were maintained in a growth chamber with a day/night temperature of 28/20°C, a 12-hour photoperiod, and medium intensity high pressure sodium and metal halide lamps. A nutrient solution containing NH4-N: 5.99 mg/L, NO3-N: 70.39 mg/L, P: 53.92 mg/L, K: 113.83 mg/L, Ca: 170.75 mg/L, Mg: 29.35 mg/L and S: 38.64 mg/L was used for irrigation. Twenty days later, soil was gently removed from the pot and all soil/roots within 1 cm of the side and bottom of the pot were discarded. The remaining soil was washed away from fine roots over a 4 mm mesh sieve. Roots were collected, washed three times in fresh 1% sodium hexametaphosphate solution (to improve soil aggregate dispersal) by shaking at 100 rpm for 15 minutes, and sonicated in a Branson 5510 water bath for 7 minutes (Kettler et al. 2001).

**Table 1.** Crops and cultivars used in this study.

| Crop (*species*) | Cultivar | Panel: |  |  |
| --- | --- | --- | --- | --- |
|  |  | General | Strawberry | Lettuce |
| **Lettuce**<br>(*Lactuca sativa*) | Green Towers | X |  | X |
|  | Salute |  |  | X |
|  | Salinas 88 |  |  | X |
|  | Bondi |  |  | X |
| **Broccoli**<br>(*Brassica oleracea*) | Marathon F1 | X |  |  |
| **Wheat**<br>(*Triticum aestivum*) | Summit | X |  |  |
| **Cilantro**<br>(*Coriandrum sativum*) | Leisure | X |  |  |
| **Spinach**<br>(*Spinacia oleracea*) | Whale F1 | X |  |  |
| **Raspberry**<br>(*Rubus idaeus*) | Heritage | X |  |  |
| **Strawberry**<br>(*Fragaria x ananassa*) | Albion | X | X | X |
|  | Monterey |  | X |  |
|  | San Andreas | X | X |  |
|  | Fronteras |  | X |  |
|  | Portola |  | X |  |
|  | Ventana |  | X |  |

Root tips (containing apical meristem) that were 15-20 mm in length were placed on Komada’s Medium (KM) agar (Komada 1975) and incubated under continuous fluorescent lighting at room temperature for seven days. Colonies similar in morphology to *F. oxysporum* f. sp. *fragariae* were subcultured and checked for identity using the PCR test described above. For each plant, 30-50 root tips were assayed and the proportion of root tips infected by the total number of root tips was determined. The proportion of infected root tips was arcsin(square root) transformed prior to statistical analysis.

### *Fusarium oxysporum* f. sp*. fragariae* population density on living plant tissues

Hygromycin-resistant *F. oxysporum* f. sp. *fragariae* strain GL1080-K was grown on 18 potato dextrose agar plates (100 mm diameter) for 14 days at room temperature under continuous fluorescent lighting. GL1080-K-colonized agar was removed from the petri dishes and blended with 500 mL sterile deionized water for 1 minute in a Waring Commercial blender (Waring Commercial Products), at which point the solution was a homogenous, viscous slurry with no agar chunks. The slurry was hand-mixed with 10 L of autoclaved fine white sand (Capitol Sand and Gravel) until homogenous and air dried for 10 days at 25°C. Inoculated sand was re-mixed daily during the drying period. Dry, infested sand was stored at 4°C for up to three weeks prior to use. Prior to each experiment, 10 L infested sand was blended with 70 L #1 Sunshine Potting Mix (Sun Gro Horticulture) to obtain an inoculum density of approximately 2 × 10^5^ colony forming units (CFUs) per gram of sand/potting mix.

Experiments using the general panel of rotation crops (Table 1) were conducted three times to estimate *F. oxysporum* f. sp*. fragariae* population density in plant tissues. Three plants (= biological replicates) were assayed for each crop type. Seeds and transplants were sown in #1 Sunshine Potting Mix and grown for 14 days in a growth chamber with the previously described settings. Seedlings were then removed from their pots (n = 3), washed to remove adhering soil, and transplanted into the infested sand/potting mix described above. Plants were returned to the growth chamber and maintained under the conditions described above for a period of six weeks.

At the end of this growth period, plants were washed and separated into the following tissue types (McCormack et al. 2015): fine roots (< 2 mm diameter, proximal to apical meristem tissues), transport roots (≥ 2 mm diameter, proximal to taproot/crown), taproot/crown cortical tissue, and taproot/crown stele tissue. Roots greater than 2 mm diameter (transport roots) were only found on strawberry and raspberry plants. Taproots (broccoli, lettuce, spinach, and cilantro) and root crowns (wheat, raspberry, and strawberry) were in all cases defined as the below-ground enlarged root tissues that are immediately basal to the stem. Stele tissues from spinach and wheat plants were too small to separate from cortical tissues. Fine roots and transport roots were collected from the center of the pot, washed three times in 1% sodium hexametaphosphate solution, and sonicated for 7 minutes in a Branson 5510 sonicator. Taproots/crowns were surface-sterilized by rinsing in 0.1% Tween 20 solution, submerging for 10 seconds in 70% ethanol, immersing in 1% sodium hypochlorite (with continuous agitation from a magnetic stir bar) for 2 minutes, and drying thoroughly (solution percentages are volume/volume). After surface disinfestation, fresh weight was recorded for each tissue type of each plant, and tissues were blended with deionized water for 5 minutes in a Waring commercial blender. The resulting slurry was spread over the surface of KM amended with hygromycin B (100 mg/L) to achieve a final detection limit of approximately 500 CFUs per gram of plant tissue. Plates were incubated at room temperature under continuous fluorescent lighting for 6-7 days prior to counting the number of colonies with GL1080-K morphology.

We used the same experimental procedure (with three independent repetitions) on two additional crop panels: a selection of *F. oxysporum* f. sp. *fragariae* resistant and susceptible strawberry cultivars, and a panel of lettuce cultivars (Table 1). For the experiments with resistant and susceptible strawberry cultivars, four plants (= biological replicates) were used and the growth chamber had day/night temperatures of 25/18°C. The lettuce cultivar experiments were conducted exactly as described for the general panel of rotation crops, but with four biological replicates and only the taproot/crown cortex and stele tissues were assayed.

### *Fusarium oxysporum* f. sp*. fragariae* population density on plant tissues buried in soil for two weeks

Two experiments were conducted to assess *F. oxysporum* f. sp*. fragariae* population density on taproot/crown tissues after two weeks of burial in soil. Initial inoculation, cultivar selection, and growth conditions were exactly as described for experiments assaying population density of GL1080-K on living tissues of the general rotation crop panel (Table 1). However, tissues were not assayed immediately after the 6-week growth period. Instead, taproots/crowns were removed from the soil, trimmed of roots longer than 2 cm, severed from above-ground biomass 2 cm above the soil line, and re-buried in the pot from which they came. Pots were irrigated with a nutrient solution (described above) as needed to maintain soil near field capacity for 14 days, and taproots/crowns were then recovered. Cortex and stele compartments could not be distinguished after the incubation period, so whole taproot/crown pieces were cleaned of adhering soil particles, surface-sterilized with 1% sodium hypochlorite, and assayed for GL1080-K population density, as described above.

### Survival of *Fusarium oxysporum* f. sp. *fragariae* in soil for 44 weeks after crop growth and plant tissue incorporation

Seeds of five rotation crops and dormant strawberry crowns were sown in potting soil (#1 Sunshine Mix, Sun Gro Horticulture) and cultivated in a growth chamber with a 12-hour photoperiod and day/night temperatures of 28/20°C for 14 days prior to the start of the experiment. Living raspberry primocanes (first year cane growth) were maintained in the same growth chamber. After 14 days, potting soil was gently washed from roots of all plants, and four plants per crop were transplanted into 2 L pots (= four biological replicates, one plant per pot). These 2 L pots contained a soil mixture that was prepared by mixing 10 L of sand infested with GL1080-K (prepared as described above), 15 L of autoclaved fine white sand (Capitol Sand and Gravel), and 63 L of soil from a strawberry field near Watsonville, CA which had been passed through a 4 mm mesh sieve. The soil/sand mixture was prepared immediately before the start of each experiment. An additional four pots were left un-planted as a “fallow” treatment. Planted and fallow pots were returned to the same growth chamber used to generate the transplants for these experiments.

Six weeks later, plants were severed at the soil line, and fresh weight of above-ground plant biomass was recorded. All plant tissues (including roots) were cut into ∼1 cm fragments and mixed evenly with the soil, which was returned to the pot from which it was removed. For the fallow treatment (no plants), soil was removed, mixed, and returned to the pot. All pots were returned to the growth chamber, left to dry for one week, and irrigated weekly thereafter with 200 mL of charcoal-filtered water per pot.

At the time of plant tissue incorporation and 12, 24, and 44 weeks post-incorporation (wpi) the soil was removed from each pot, thoroughly mixed by hand, and approximately 150 grams of soil (excluding large tissue fragments) was retrieved for quantification of GL1080-K population density. The abundance of GL1080-K in the soil was quantified using soil dilution plating on KM amended with hygromycin B (100 mg/L). Two technical replicates were conducted per biological replicate at each timepoint and the average of these two technical replicates used in statistical analyses.

At the final timepoint (44 wpi), soil was passed through a 4 mm mesh sieve and plant tissue fragments larger than 4 mm were collected and dried at 30°C for 24 hours. Tweezers were used to scrape large adhering soil particles from the surface of tissues, which were not otherwise cleaned. Strawberry tissues were separated into transport roots and crown tissues based on clear morphological differences (Supplementary Fig. S1). Tissues were weighed and ground into a fine powder with a mortar and pestle. This powder was suspended in 1% sodium hexametaphosphate and dispensed onto KM amended with 100 mg hygromycin B per L (detection limit was between 8 and 1,000 CFU/g).

### Statistical analysis

All statistical analyses were conducted in R (version 3.4.3) (R Core Team 2013). Levene’s test (“leveneTest”, package = “car”) was used to assess homogeneity of variance (Fox and Weisberg 2011), and normality was checked by generating quantile-quantile plots (“qqnorm”, package = “stats”). For data with a continuous response variable, linear mixed-effects models were fit by “lmer” from the “lmerTest” package (Kuznetsova et al. 2017). For the full mixed model, crop type was coded as a fixed effect, while experiment and experiment by crop interactions were coded as random effects. Chi-squared tests were conducted between the full model and models where one random effect was removed to test the significance of the removed variables (“anova” with “type=Chisq”, package = “lmerTest”). Two-way ANOVAs were conducted with “anova” from the “lmerTest” package. Multiple tissue types from a single plant or separate timepoints in the soil survival experiments were not considered independent variables, so separate two-way ANOVAs were conducted for each tissue type or timepoint. Paired and Welches t-tests were conducted with “t.test” (package = “stats”), and “p.adjust” with “method = bonferroni” used to correct for multiple comparisons. Tukey’s honestly significant difference (HSD) tests for multiple comparisons were conducted with “HSD.test” from the “agricolae” package (α = 0.05) (de Mendiburu 2017). “cor.test” (package = “stats”) was used to test the significance of correlation (method = “pearson”). *R*^2^ values were calculated by “lm” (package = “stats”).

## RESULTS

### *Agrobacterium tumefaciens*-mediated transformation of *F. oxysporum* f. sp. *fragariae*

Co-incubation of fungal spores and *A. tumefaciens* cells led to the emergence of 2-5 hygromycin B-resistant colonies of *F. oxysporum* f. sp. *fragariae* per plate. PCR products of the expected size (∼1.0 kb) were obtained from twelve transformants using HPH primers (specific for the hygromycin gene, *hph*). Six of those twelve transformants were confirmed to have retained pathogenicity on the strawberry cultivar, Albion, and to have growth rates on PDA and KM which were indistinguishable from that of the wild type isolate (data not shown). A single transformant resulting from this protocol, called “GL1080-K”, was used in subsequent experiments.

### Root tip colonization in naturally infested field soil

All crops tested in these experiments sustained root tip infections by *F. oxysporum* f. sp *fragariae*. The highest rates of root infection were observed on Fusarium wilt*-*susceptible strawberry cultivar (Albion), resistant strawberry cultivar (San Andreas), and raspberry (Fig. 1). The lowest root infection rates were observed on broccoli and spinach plants (Fig. 1). A two-way ANOVA revealed a significant effect of crop on the frequency of root tip infection (*P* < 0.001), whereas experiment and crop by experiment interaction were not significant (*P* = 0.183 and *P* > 0.999, respectively). Mean inoculum densities of *F. oxysporum* f. sp*. fragariae* (± standard error) were 380 (±127), 260 (±70), and 540 (±42) at the start of experiments one, two, and three, respectively. Soil inoculum density was positively correlated with the frequency of root infection (total infected roots / total roots assayed) across the three experiments (n = 3, R^2^ = 0.99, *P* = 0.023, data not shown).

**Fig. 1.**
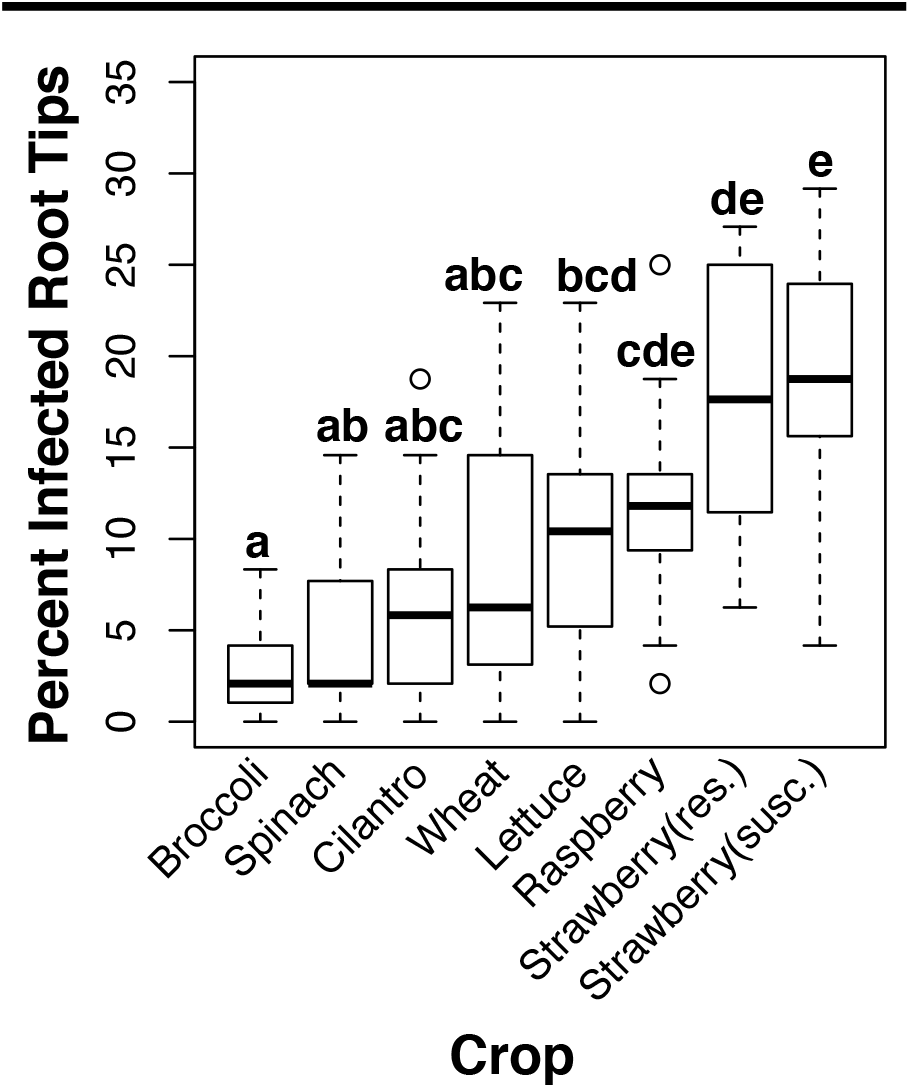
Box plot depicting the rate of root tip infection by *Fusarium oxysporum* f. sp. *fragariae* in naturally infested soil. Strawberry cultivars San Andreas and Albion are resistant and susceptible to Fusarium wilt, respectively. Tukey’s honest significant difference statistical groupings were determined from arcsin(squared root) transformed data.

### *Fusarium oxysporum* f. sp*. fragariae* population density on living plant tissues

#### General rotation crop panel

After six weeks of growth in soil infested with *F. oxysporum* f. sp. *fragariae* (isolate GL1080-K), the pathogen was detected on the taproot/crowns and/or fine roots of all tested rotation crops (Table 2). Raspberry transport roots contained the highest mean population of any living tissue from a non-strawberry crop, and these were not significantly different from transport roots of susceptible strawberry (Table 2). Logarithmic (base 10) transformation was conducted on CFU counts +1 to achieve a log-normal distribution. Two-way ANOVAs from crown cortex, crown stele, and fine roots revealed a highly significant effect of crop (*P* ≤ 0.006) and no significant effects of experiment or crop by experiment interaction (*P* ≥ 0.711).

**Table 2:** Colonization of living rotation crop tissues by *Fusarium oxysporum* f. sp. *fragariae* (strain GL1080-K).

| Crop<br>(Cultivar) | Fine Root<sup>z</sup> | Transport Root<sup>z</sup> | Taproot/crown |  |  |
| --- | --- | --- | --- | --- | --- |
|  |  |  | Cortex<sup>z</sup> | Stele<sup>z</sup> | Total<sup>z,y</sup> |
| **Lettuce**<br>(Green Towers) | 0 (0.0) b | NT | 0.27 (0.25) a | 0 (0.0) a | 0.24 (0.24) a |
| **Spinach**<br>(Whale) | 1.74 (0.41) ab | NT | NT | NT | 0.62 (0.41) a |
| **Broccoli**<br>(Marathon F1) | 0.84 (0.4) ab | NT | 0.27 (0.25) a | 0.24 (0.22) a | 0.25 (0.24) a |
| **Cilantro**<br>(Leisure) | 1.92 (0.34) a | NT | 0 (0.0) a | 0 (0.0) a | 0 (0) a |
| **Wheat**<br>(Summit) | 0.85 (0.4) ab | NT | NT | NT | 0.3 (0.3) a |
| **Raspberry**<br>(Heritage) | 0.52 (0.35) ab | 3.15 (0.45) a | 0.55 (0.35) a | 0.7 (0.34) a | 0.84 (0.36) a |
| **Strawberry**<br>(San Andreas) | 1.42 (0.54) ab | 2.57 (0.76) a | 3.09 (0.43) b | 0.24 (0.23) a | 2.78 (0.44) b |
| **Strawberry**<br>(Albion) | 0.27 (0.25) ab | 2.93 (0.4) a | 4.35 (0.33) b | 5.04 (0.21) b | 4.93 (0.23) c |

<sup>z</sup>Data are presented as: mean log<sub>10</sub>(CFUs + 1/g), followed by standard error (in parentheses). Means in the same column followed by the same letter are not significantly different ( $\alpha = 0.05$ ).
<sup>y</sup>Values for total taproot/crown tissues for crops whose tissues were separated into cortex/stele represent the weighted average of log<sub>10</sub>(CFUs + 1/g) from these tissues (i.e. log<sub>10</sub>(CFUs + 1/g) normalized to the observed weight).
NT: tissue type was not tested.

Fine roots appear to be an important site of inoculum production on lettuce, spinach, broccoli, cilantro, and wheat. For example, population densities of GL1080-K on cilantro fine roots were significantly higher than cilantro taproot cortex (paired t-test, Bonferroni corrected *P* < 0.001). In contrast, fine root population densities were significantly lower than populations in crown cortex tissues of both strawberry cultivars (paired t-test with Bonferroni correction; cv. San Andreas *P* = 0.039, cv. Albion *P* < 0.001). This suggests that fine roots are less important for production of soilborne inoculum on strawberry and raspberry plants, because greater population densities are produced on other tissues.

#### Resistant/susceptible strawberry cultivars

After 14 days of growth in infested soil, population densities of GL1080-K were determined for transport roots, crown cortices, crown steles, and petiole tissues of all six cultivars (Fig. 2A). Two-way ANOVAs revealed cultivar to be a significant source of variation for all tissue types (*P* < 0.001), whereas experiment and crop by experiment interaction were not significant (*P* ≥ 0.321). Tukey’s HSD showed population densities of GL1080-K to be significantly lower in resistant cultivars than in susceptible cultivars for all tissue types (Fig. 2B). Both susceptible cultivars, Albion and Monterey, had higher densities of *F. oxysporum* f. sp*. fragariae* in stelar tissues than were observed in cortices. The difference was significant for Albion (*P* = 0.036) but not Monterey (*P* = 0.101) (Fig. 2B). In contrast, all resistant cultivars had significantly lower population densities in the stele than in the cortex (*P* ≤ 0.002) (Fig. 2B).

**Fig. 2.**
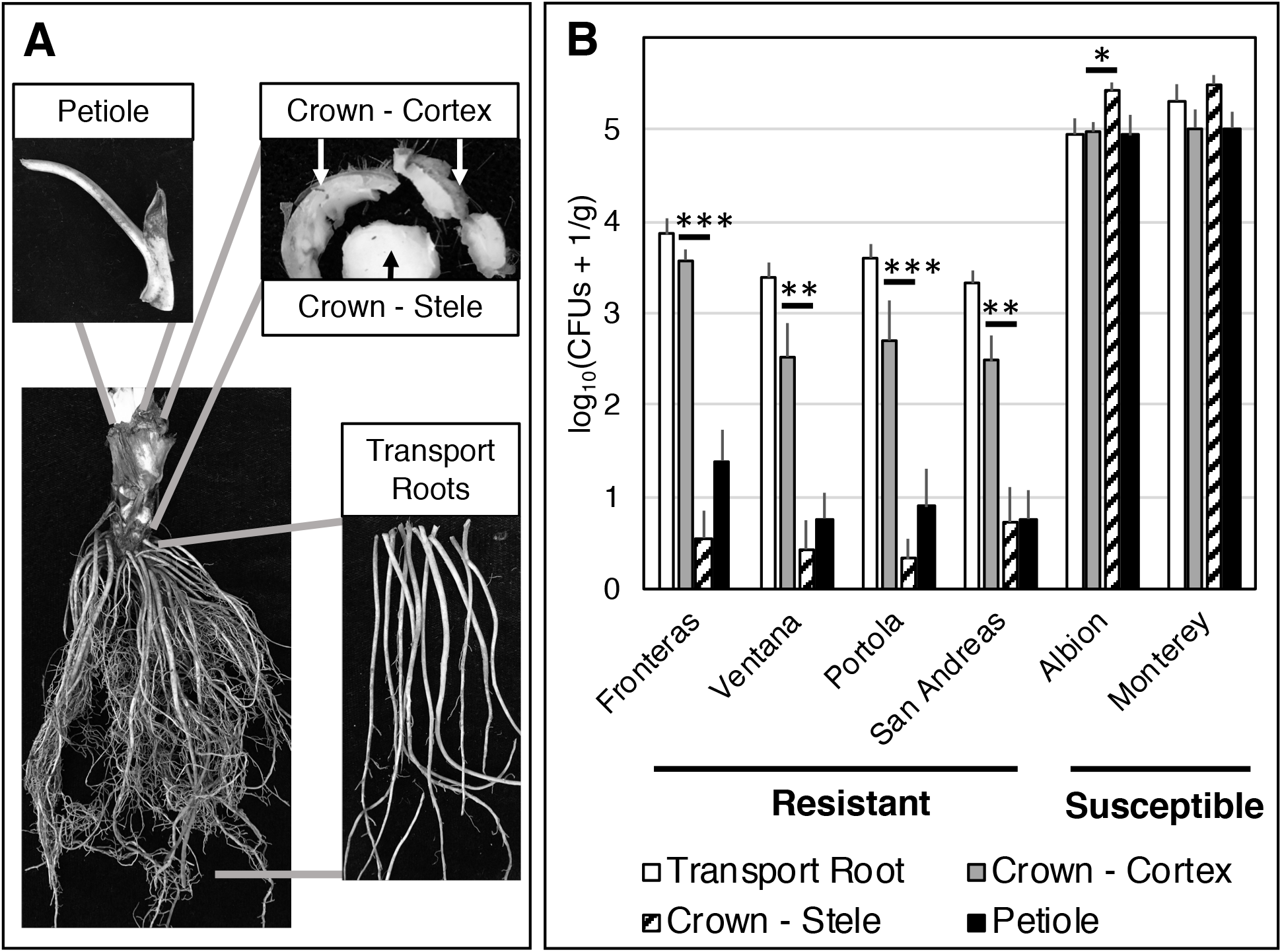
Colonization of resistant and susceptible strawberry cultivars by *Fusarium oxysporum* f. sp. *fragariae* (GL1080-K). **A.** Four tissue types that were assayed. **B.** Mean log10(CFUs of GL1080-K + 1/g) recovered from the assayed tissues. Error bars denote standard error of the mean. For all tissue types, the only statistically significant difference was between resistant and susceptible cultivars (*** = P < 0.001, ** = P < 0.01, * = P < 0.05).

#### Lettuce cultivar panel

GL1080-K population densities were quantified from the taproot cortex and stele tissues of four lettuce cultivars differing in susceptibility to Fusarium wilt of lettuce, and the susceptible strawberry cultivar, Albion. *Fusarium oxysporum* f. sp. *fragariae* was not detected in either the cortex or the stele of the lettuce cultivar Salinas 88 but was recovered from the stele of all other lettuce cultivars (Table 3), including two cultivars with resistance to Fusarium wilt of lettuce (caused by *F. oxysporum* f. sp. *lactucae*). For both cortical and stelar tissues, two-way ANOVA revealed plant genotype to be a significant source of variation in GL1080-K population densities, reflecting significantly higher levels in strawberry than in any of the lettuce cultivars (Table 3).

**Table 3:** Colonization of lettuce taproot tissues by *Fusarium oxysporum* f. sp. *fragariae*.

| Crop (cultivar) | Leaf Habit | Susceptibility<sup>z</sup> | Cortex<sup>y,w</sup> | Stele<sup>x,w</sup> |
| --- | --- | --- | --- | --- |
| **Lettuce** (Bondi) | Romaine | Susceptible | 0.21 (0.21) a | 0.49 (0.25) a |
| **Lettuce** (Green Towers) | Romaine | Resistant | 0 (0) a | 0.14 (0.14) a |
| **Lettuce** (Salinas 88) | Head | Susceptible | 0 (0) a | 0 (0) a |
| **Lettuce** (Salute) | Head | Resistant | 0 (0) a | 0.49 (0.33) a |
| **Strawberry** (Albion) | N/A | N/A | 4.41 (0.43) b | 5.55 (0.08) b |

<sup>z</sup>Lettuce cultivars were either resistant or susceptible to *Fusarium* wilt of lettuce, caused
by *Fusarium oxysporum* f. sp. *lactucae*.
<sup>y</sup>Cortex of the taproot was assayed separately from the stele.
<sup>x</sup>Stele of the taproot was assayed separately from the cortex.
<sup>w</sup>Data are presented as mean log<sub>10</sub>(CFUs + 1/g of plant tissue), standard error (in
parentheses). Means in the same column followed by the same letter are not
statistically different ( $\alpha = 0.05$ ). The average limit of detection in these assays was 187
CFUs/g.

### *Fusarium oxysporum* f. sp*. fragariae* population density on plant tissues buried in soil for two weeks

Taproot/crown tissues recovered two-weeks post burial were not separated into cortex and stele (as living tissues were), so total GL1080-K CFUs/g in living taproot/crown tissues were calculated from the weighted average of cortex and stele population densities (Table 2). Welch’s two-sample t-test was used to evaluate differences in log10(GL1080-K CFUs +1/g) between living and post-burial taproots/crowns for each crop, with Bonferroni correction for multiple comparisons applied to all *P-*values (Fig. 3). GL1080-K population densities were significantly greater after taproot/crown burial for broccoli, spinach, cilantro, and wheat (*P* = 0.002, *P* = 0.015, *P* = 0.033, and *P* = 0.008, respectively), but not for either strawberry cultivar, raspberry, or lettuce (*P* = 0.324 for raspberry, *P* > 0.999 for both strawberry cultivars and lettuce) (Fig. 3).

**Fig. 3.**
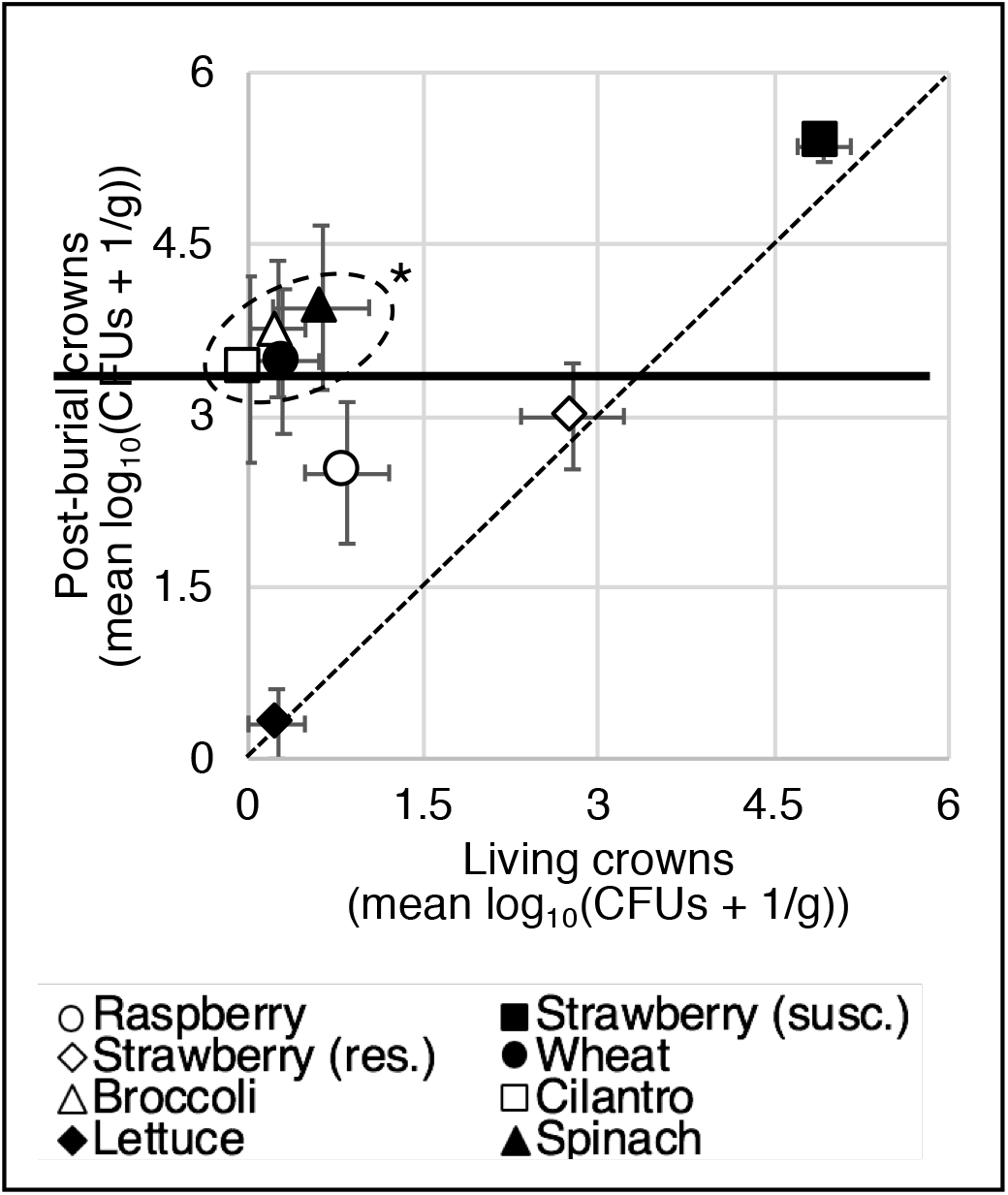
*Fusarium oxysporum* f.sp. *fragariae* (GL1080-K) population density on living and two-weeks post-burial taproots/crowns. Mean log_10_(CFUs + 1/g) of GL1080-K from total living crown tissues are shown on the x-axis, and post-burial GL1080-K population densities are on the y-axis. Error bars denote standard error of the mean. Significantly higher post-burial GL1080-K populations are denoted by: * = P < 0.05. Strawberry cultivars were either resistant (cv. San Andreas) or susceptible (cv. Albion) to Fusarium wilt of strawberry.

### Survival of *Fusarium oxysporum* f. sp. *fragariae* in soil for 44 weeks after crop growth and plant tissue incorporation

#### Soil inoculum density

Experiment, but not crop treatment, had a significant effect on log_2_(GL1080-K CFUs/g of soil) at the time of plant tissue incorporation (experiment *P* = 0.002; crop *P* = 0.155; crop by experiment interaction *P* = 0.896; two-way ANOVA with crop as a fixed effect, experiment as a random effect, and interaction as a random effect). Total variance increased significantly between the time of incorporation and 12 wpi (Supplementary Fig. S2).

Two-way ANOVA with the linear mixed model described above showed a significant crop effect for soil population densities at 12, 24, and 44 wpi (*P* < 0.001), and a significant experiment effect only at 12 wpi (*P* < 0.001). A repeated measures ANOVA was conducted to assess the significance of time on soil population densities of GL1080-K. Crop, time, and crop by time interactions were considered fixed effects, and all were highly significant (*P* < 0.001).

Crop treatments were compared based on changes in soil inoculum densities of GL1080-K that occurred between plant incorporation and later timepoints. To accomplish this analysis, differences (i.e. ∆log_2_(GL1080-K CFUs/g)) were calculated between population densities at 12, 24, and 44 wpi and the time of incorporation (0 wpi) for each pot (biological replicate) (Fig. 4). Two-way ANOVAs with ∆log_2_(GL1080-K CFUs/g) as the dependent variable, crop as a fixed effect, and experiment and crop by experiment interaction as random effects were conducted for each timepoint. These analyses revealed a significant experiment effect only at 12 wpi (*P* < 0.001), but all other random effects from all timepoints were not significant (*P* > 0.999). Tukey’s HSD groups were calculated for 24 and 44 wpi (Supplementary Table S1). These analyses revealed that, for both timepoints, soil in pots where raspberry was grown had significantly higher inoculum densities than soil in pots in which other crops were grown. For lettuce, broccoli, and spinach, inoculum densities at 24 wpi were not significantly different from the fallow (no plant) treatment.

**Fig. 4.**
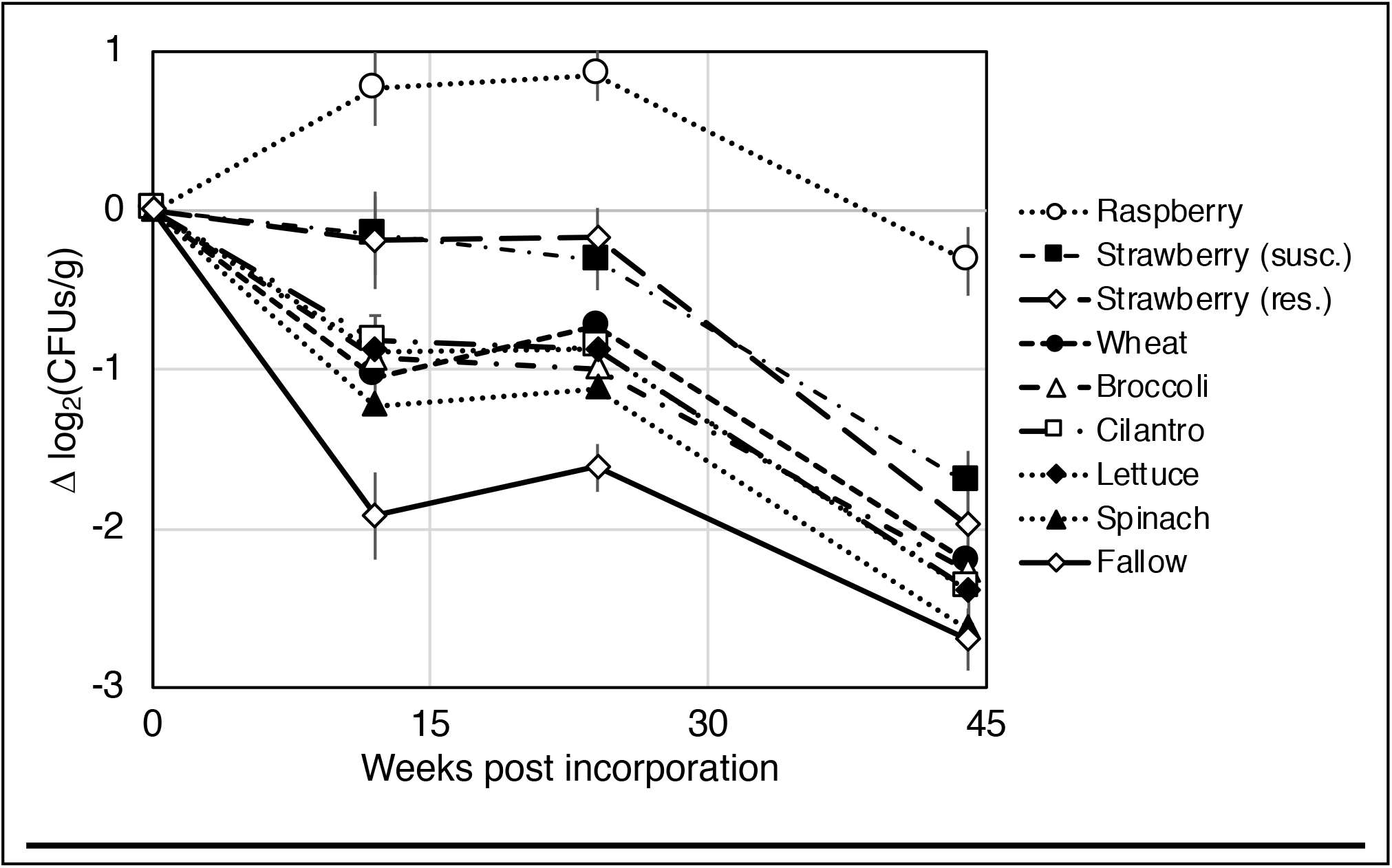
Changes in mean soil population densities of *Fusarium oxysporum* f. sp. *fragariae* (GL1080-K) after crop debris incorporation. Differences (i.e. Δlog_2_(CFUs/g) were calculated between GL1080-K CFUs/g of soil at 12, 24, and 44 weeks post incorporation and the time of crop debris incorporation. Error bars denote standard error of the mean. Strawberry cultivars were either resistant (cv. San Andreas) or susceptible (cv. Albion) to Fusarium wilt of strawberry.

#### Recovery of plant tissue fragments at 44 weeks post incorporation

Crop was a highly significant (*P* < 0.001; two-way ANOVA) source of variation in weight of plant tissues recovered at 44 wpi (*P* > 0.999 for random effects: experiment and crop by experiment interaction). Significantly more tissue was recovered from perennial crops (raspberry and strawberry) compared to annual crops (Table 4), and this rate of recovery was not correlated with fresh weight of plant tissues at the time of incorporation (data not shown).

**Table 4.**
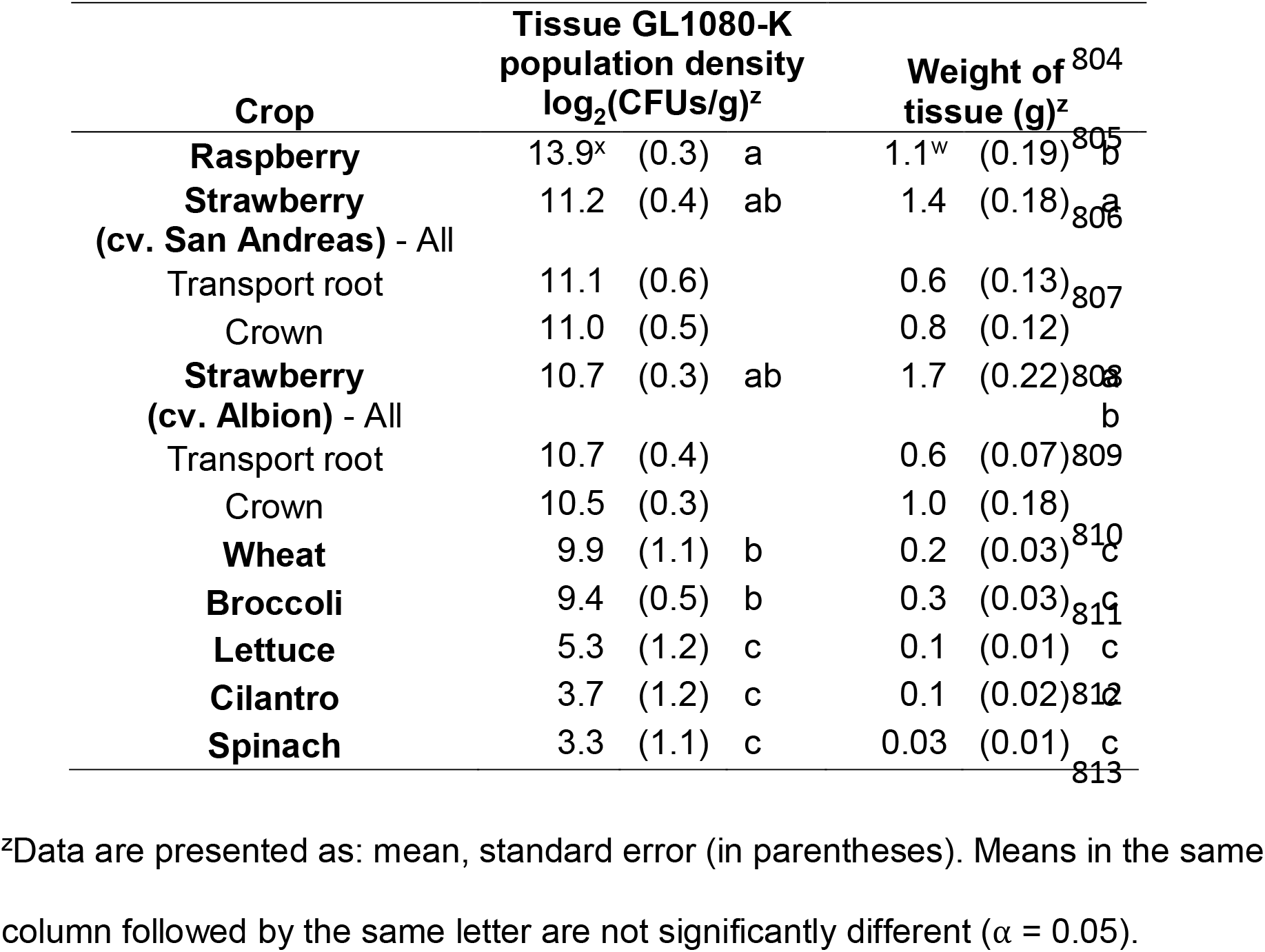
Weight of, and *Fusarium oxysporum* f. sp*. fragariae* (GL1080-K) population densities on, plant tissues recovered at 44 weeks post plant incorporation into soil.

#### Population densities of GL1080-K on recovered tissue fragments

Two-way ANOVAs conducted on the GL1080-K population density (log_2_(CFUs + 1/g)) in tissues recovered at 44 wpi also revealed a highly significant crop effect (*P* < 0.001), and no significant random effect terms (*P* > 0.999 for both experiment and crop by experiment interaction) (Table 4).

Soil and plant tissue population densities of GL1080-K (44 wpi) were compared within treatments by paired t-tests. Significantly higher population densities of GL1080-K were observed in tissues from raspberry and both strawberry cultivars compared with soil populations in their corresponding pots (*P* < 0.001). For spinach and cilantro, population densities of GL1080-K were significantly lower in tissue than in soil (*P* = 0.005 and *P* = 0.032, respectively). The difference between tissue and soil was not significant for the wheat, lettuce or broccoli treatments (*P* > 0.999, *P* = 0.212, and *P* = 0.500).

A weak, but significant, positive correlation was observed between CFUs of GL1080-K per gram of tissue fragments recovered at 44 wpi and mean soil CFUs/g from 12, 24, and 44 wpi (n = 78). Biological replicates from which tissue fragments were not recovered were excluded from this analysis. *R*^2^ values for 12, 24, and 44 wpi were 0.31, 0.37, and 0.36 respectively, all of which were highly significant (*P* < 0.001)

(Supplementary Fig. S3). The *R*^2^ value was 0.43 for tissue CFUs/g vs. soil CFUs/g at 44 wpi when extreme outliers (i.e. points that were more than 4.5 times the interquartile range; n=72) were excluded (Fig. 5).

**Fig. 5.**
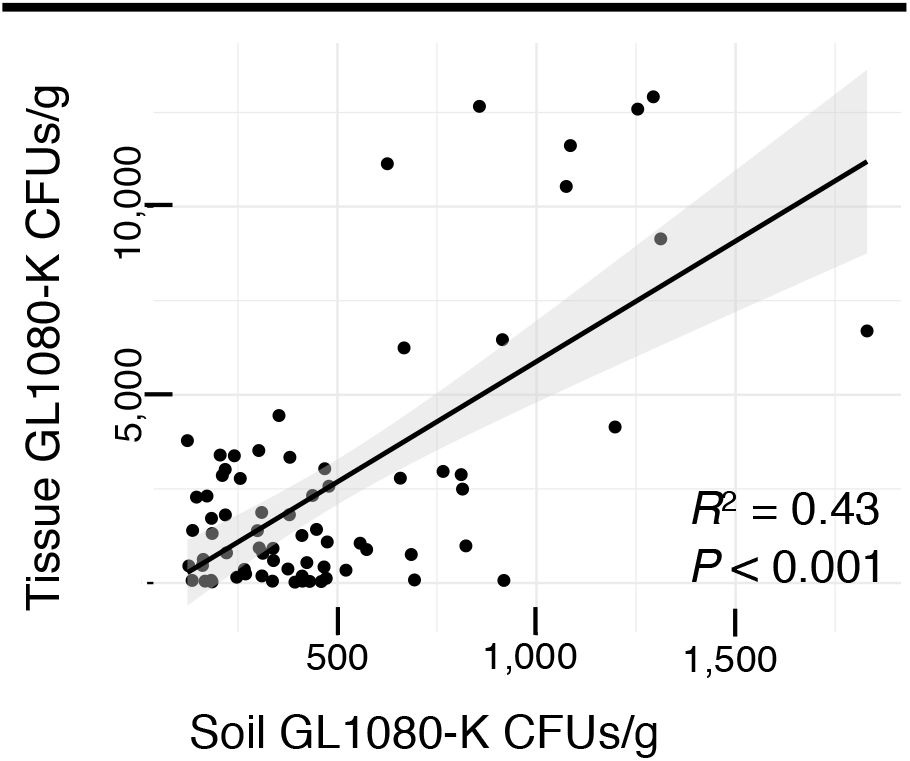
Correlation between *Fusarium oxysorum* f. sp. *fragariae* (GL1080-K) CFUs per gram from soil and tissues recovered at 44 weeks post inoculation. Pots (biological replicates) from which values were beyond 4.5 times the interquartile range for either soil or tissue CFUs/g were excluded from this analysis (n = 72).

## DISCUSSION

This study shows, for the first time, a clear relationship between reproduction on asymptomatic hosts and long-term soil persistence for a Fusarium wilt pathogen. All crops and cultivars tested could be infected by *F. oxysporum* f. sp. *fragariae*, but significant differences were observed in the extent of reproduction in their tissues. Annual crops (broccoli, lettuce, spinach, wheat, and cilantro) supported the lowest population densities of *F. oxysporum* f. sp*. fragariae*. Raspberry and Fusarium wilt-resistant strawberry cultivars supported populations and infection rates that were consistently closest to those observed on a susceptible strawberry cultivar. Population densities on asymptomatic hosts were typically greater after residue incorporation, highlighting the need to consider this phase of pathogen growth and reproduction in evaluating potential rotation crops.

Long-term experiments in infested fields directly assess the utility of crop rotations, but such sites are not always available for research. In their absence, the methods used in our soil survival experiments provide a useful template for assessing inoculum production on asymptomatic hosts. Artificially infesting field soil with an antibiotic selectable transformant (in controlled environments) enables rapid quantification yet maintains a biological environment that is similar to what would exist in a commercial field. Assaying soil populations at multiple timepoints and tissue populations after a long incubation period (such as 44 weeks) can provide a useful indication of the plant’s total inoculum contribution to soil. Soilborne populations observed by this procedure reflect total growth from rhizosphere, saprophytic activity in soil, and growth after decomposition of all tissue types. Plant species and tissues decompose at different rates (Table 4), so assaying tissues at the end of the incubation period is necessary to reveal inoculum that has not yet been released into the soil. For these reasons, the soil survival experiments described in this study are likely to be straightforward and effective for identifying crops that pose a risk for pathogen persistence.

Soil survival experiments showed that inoculum produced on plant tissues had a long-term impact on *F. oxysporum* f. sp. *fragariae* populations. The type of crop had little impact on bulk soil populations of GL1080-K prior to incorporation. In contrast, total variance increased significantly after plant residue incorporation. This shows that differences in bulk soil GL1080-K populations between crops were associated with decomposition of plant tissues. A positive correlation (*R*^2^ = 0.43, *P* < 0.001) was observed between soil and tissue populations of *F. oxysporum* f. sp*. fragariae* at 44 wpi. The fallow (no-plant) treatment consistently had the lowest mean inoculum density at all timepoints post-incorporation, which is consistent with observations that all crops can support inoculum production.

Data on the colonization of living plant tissues by *F. oxysporum* f. sp*. fragariae* suggest that raspberry would pose a greater threat than other non-strawberry crops for maintaining soilborne populations of this pathogen (Fig. 1; Table 1). The magnitude of this threat is much greater when the consequences of residue incorporation are taken into account. The density of *F. oxysporum* f. sp*. fragariae* CFUs/g in raspberry tissue fragments at 44 wpi was more than 10 times greater than what was observed in the most densely colonized living raspberry tissues (Table 2; Table 4). This indicates that pathogen exploitation of tissue following incorporation into soil can greatly amplify the amount of inoculum added to the soil population. This observation suggests that any evaluation of rotation crops should include assays of pathogen populations after crop residue incorporation.

The *F. oxysporum* f. sp. *fragariae-*resistant cultivars tested in our experiments are heterozygous for a single, dominant resistance gene: *Fw1* (Pincot et al. 2018). In these cultivars, significantly lower inoculum densities were observed in the vascular stele than in cortical tissues (Fig. 2B). This suggests that *Fw1*-mediated resistance limits access to vascular tissues but does not prevent colonization of cortical tissues. These results are consistent with previous reports showing that pathogenic strains of *F. oxysporum* can extensively colonize resistant cultivars of their preferred hosts (Elmer and Lacy 1987; Gordon et al. 1989; Houterman et al. 2009; Rishbeth 1955; Scott et al. 2014; van der Does et al. 2018). What are the genetic determinants that enable *F. oxysporum* f. sp. *fragariae* to colonize resistant cultivar cortical tissues at a significantly greater level than other crops (Fig. 2B)? Do the same factors enable colonization of raspberry? Such questions have received little attention in the literature but have significant implications for our understanding of host specificity and the persistence of Fusarium wilt pathogens in the absence of their preferred host.

It is intriguing to note that raspberry, the most extensively colonized rotation crop, is more closely related to strawberry than the other crops tested. Consistent with our findings, previous studies showed other Fusarium wilt pathogens to reproduce more extensively on plants that are in the same family as their preferred host (Dhingra and Coelho Netto 2001; Haware and Nene 1982). Furthermore, a wilt disease of raspberry caused by *F. oxysporum* was recently reported in Peru, and the causal agent was also pathogenic to strawberry (Martin et al. 2017). Likewise, a recently described pathogen of blackberry, *F. oxysporum* f. sp. *mori*, is also pathogenic to strawberry (Pastrana et al. 2017b). These observations suggest there are common genetic determinants of susceptibility to *F. oxysporum* in raspberry, blackberry and strawberry. Analysis of asymptomatic plant colonization in plant species other than the preferred host may provide insight into the prerequisites for host specific pathogenicity in the *F. oxysporum* species complex.

Whereas soil survival experiments can assess total inoculum contribution to soil, data from infection of living plant tissues can provide information about where this inoculum is being produced. For example, our data suggest that localized fine root infections serve as an important site of *F. oxysporum* f. sp*. fragariae* population growth on some asymptomatic hosts, but not on the susceptible strawberry cultivar, Albion. Fine roots have a greater surface area to volume ratio than crowns and are therefore more likely to come in proximity of soilborne propagules of *F. oxysporum* f. sp*. fragariae*. If infections remain localized, many fine root infections, resulting from a 5-20% rate of root infection (Fig. 1), would be expected to contribute more to soilborne inoculum than a limited number of localized crown infections. In contrast, the greatest population densities on the susceptible strawberry cultivar, Albion, occurred on crown stele tissues, which reflects more extensive colonization of these tissues compared to fine roots. GL1080-K populations on Albion fine roots were not significantly different from those on asymptomatic hosts (Table 2) and were significantly lower than any other Albion tissue type (*P* < 0.005). Diverse root structure and function can exist within fine roots defined by diameter (McCormack et al. 2015), and future research may uncover additional patterns of colonization that are specific to root function.

Past research has shown that fields can remain infested for decades by Fusarium wilt pathogens in the absence of a susceptible host (Rishbeth 1955; Smith et al. 2001). However, the rate of soilborne pathogen population decline observed here and in other studies (Banihashemi and Dezeeuw 1973; Elmer and Lacy 1987; Gordon and Koike 2015; Rishbeth 1955; Scott et al. 2012; Stover 1956; Vakalounakis and Chialkas 2004), suggest that few chlamydospores survive long enough to account for pathogen persistence over many years. Rather, it seems likely that persistence of Fusarium wilt pathogens is attributable to their activity as parasites of non-susceptible crops and their ability to further colonize host tissue as it decomposes.

Long-term survival in soil may be enhanced when pathogen propagules are embedded in recalcitrant plant tissues. Crop debris from perennial plants, such as strawberry and raspberry, were recovered at significantly higher rates than annual crops (44 wpi), and recovery rates were not correlated with plant fresh weight at the time of incorporation (data not shown). The density of pathogen populations on these tissues averaged 4-12 times higher than that of the surrounding soil (Table 4). Thus, tissues that are slow to degrade may extend the interval during which a pathogen is protected from attrition in the bulk soil environment, thereby increasing the risk for persistence associated with perennial crops.

The length of rotation required before a susceptible host can be grown depends on many factors, including the threshold of inoculum required to cause disease. Rishbeth (1955) observed that *F. oxysporum* f. sp. *cubense* was typically undetectable within 8 years after a banana crop, and yet this interval did not prevent development of disease in susceptible bananas. The assay procedure used by Rishbeth (1955) had an estimated detection threshold of 2 propagules per gram. Such a low density of inoculum may be sufficient to cause disease in perennial crops, such as banana, because the probability of rare infection events will increase as roots explore the soil over time. In contrast, 10 CFUs of *F. oxysporum* f. sp. *lactucae* per gram would be unlikely to cause economically damaging losses in lettuce, a 65 to 80 day crop, even on a susceptible cultivar (Gordon and Koike 2015).

The inoculum threshold for observing disease by *F. oxysporum* f. sp. *fragariae* on strawberries has not been characterized, but 50-70 CFUs/g have been observed in heavily diseased fields (Henry, unpublished data). The threshold for disease may be quite low for strawberries, given that plantings are maintained for 6-12 months. Consequently, crop rotation should be supplemented by other disease management strategies, such as cultivar selection.

Core findings from this study are congruent with past observations and likely to be broadly relevant in Fusarium wilt pathosystems. Most crops are expected to support asymptomatic infection by Fusarium wilt pathogens, but crops may differ in the extent to which they become colonized. *Fusarium oxysporum* is likely to grow more extensively after incorporation of crop residue, indicating that assays of only living plant tissues may not provide a complete picture of the impact on pathogen soil populations. Infection of fine roots appears to be the rule for Fusarium wilt pathogens, and presumably is a mechanism by which persistence can occur for extended periods in the absence of a susceptible host. However, if infections remain localized, fine root colonization would not be expected to prevent a significant decline in soil inoculum density over time (Gordon 2017). On the other hand, the more extensive colonization that can occur in transport roots and crown/taproot tissues could significantly reduce the anticipated benefit of crop rotation. Resistant cultivars and crops that are closely related to the susceptible host may pose a greater risk for pathogen population growth and persistence in soil. Future controlled-environment experiments designed to assess the extent to which asymptomatic hosts support growth of Fusarium wilt pathogens should include assays of host tissues and soil after incorporation of crop residue.

## ACKNOWLEDGMENTS

We thank our funding sources for supporting this research: the Storkan-Hanes-McCaslin Foundation, the Western Sustainable Agricultural Research and Education Program (WSARE; Graduate Student Grant GW17-032), the California Strawberry Commission, USDA-NIFA through Hatch project CA-D-PPA-6699-H, and the National Science Foundation Graduate Research Fellowship Program. We also thank Melodie Najarro, Lia Lopez, Mariel Munji, Samuel Koehler, Megan Haugland, Madeira Alba, and Bradley Jenner for technical assistance with experiments.

---

**Supplementary Figure S1.** Representative photos of tissues obtained after 44 weeks post plant incorporation corresponding to the following crops/tissues: A. strawberry crown tissues, B. strawberry transport roots, C. raspberry, D. broccoli, E. lettuce, F. wheat, G. spinach, and H. cilantro.

**Supplementary Figure S2.** Soil populations of *Fusarium oxysporum* f. sp. *fragariae* (log_2_(CFUs/g)) observed at the time of plant incorporation and 12, 24, and 44 weeks post incorporation. Data are presented from experiments one, two, and three separately (panels A, B, and C, respectively).

**Supplementary Figure S3.** Correlation between *Fusarium oxysorum* f. sp. *fragariae* GL1080-K CFUs per gram from soil and tissues recovered at 44 weeks post inoculation. All pots (biological replicates) from which tissues were recovered are included in this analysis (n = 78).

